# Random electrical noise drives non-deterministic computation in cortical neural networks

**DOI:** 10.1101/2022.12.03.518981

**Authors:** Elizabeth A Stoll

## Abstract

In cortical neurons, spontaneous membrane potential fluctuations affect the likelihood of firing an action potential. Yet despite retaining sensitivity to random electrical noise in gating signaling outcomes, these cells achieve highly accurate computations with extraordinary energy efficiency. A new approach models the inherently probabilistic nature of cortical neuron firing as a thermodynamic process of non-deterministic computation. Typically, the cortical neuron is modeled as a binary computational unit, in either an off-state or an on-state, but here, the cortical neuron is modeled as a two-state quantum system, with some probability of switching from an off-state to an on-state. This approach explicitly takes into account the contribution of random electrical noise in gating signaling outcomes, particularly during cortical up-states. In this model, the membrane potential is described as the mixed sum of all component microstates, or the quantity of von Neumann entropy encoded by the computational unit. This distribution of macrostates is given by a density matrix, which undergoes a unitary change of basis as each unit, ‘System A’, interacts with its surrounding environment, ‘System B’. Any linear correlations reduce the number of distinguishable pure states, leading to the selection of an optimal system state in the present context. This process of information compression is shown to be equivalent to the extraction of predictive value from a thermodynamic quantity of information. Calculations demonstrate that estimated coulomb scattering profiles and decoherence timescales in cortical neurons are consistent with a quantum system, with random electrical noise driving signaling outcomes.

## I. INTRODUCTION

Mammalian spinal reflex circuits reliably transmit information, with signaling outcomes that are easily predicted by analyzing upstream activity [1–3]. Meanwhile, cortical neuron firing patterns are highly unpredictable [4], with stochastic ion leak and spontaneous subthreshold fluctuations in membrane potential significantly contributing to the likelihood of firing an action potential [5, 6]. Unlike spinal reflex circuits, which are robust to this kind of random electrical noise, cortical neurons actively manage excitation and inhibition to achieve a coordinated ‘up-state’, hovering near action potential threshold and allowing random noisy events to drive signaling outcomes [7]. A major challenge in modern neuroscience is understanding how noisy coding at the synaptic level can be reconciled with the accurate and energy-efficient coding that is observed at the neuronal population level.

The Hodgkin-Huxley equations provide a good approximation for predicting cortical neuron firing patterns under steady-state conditions [8]. Repetitive firing patterns even emerge in this model when differentiating with respect to time and applied current density [9, 10]. Linear approximations of the Hodgkin-Huxley model also accurately predict shifts in membrane potential, as long as temperatures are below 27°C and the region of membrane being modeled is above 200 square microns in size [11]. But it is worth noting that the underlying relationship between membrane voltage, ion conductances, and channel activation – given by these four partial differential equations – must ultimately be described by either modeling all eigenvectors in the system along real and complex planes, or by modeling a Hopf bifurcation to find the critical point where the cortical neuron flips from an off-state to an on-state [12–14]. Both of these deeper models essentially describe quantum processes, utilizing imaginary axes to approximate the contribution of inherently random events to signaling outcomes.

In the classical view, neural computations are encoded by the relative timing or rate of action potentials [15, 16]. With this approach, the neuron is essentially modeled as a transistor – always in an on-state or an off-state, spiking or not spiking. A new model, presented here, formalizes the inherently probabilistic nature of cortical neuron firing, with each cell computing the probability of transitioning from an off-state to an on-state. Using this approach, the cortical neuron encodes far more information than its Shannon entropy; it encodes von Neumann entropy, or the mixed sum of all component microstates. Here, the cortical neuron is a two-state quantum system, with some *probability* of firing an action potential. Since the present voltage state of the macro-scale computational unit is dependent on inherently random events, this method models the state of a cortical neuron as the mixed sum of all component microstates, or the physical quantity of information held by the computational unit.

Here, the probability of a cortical neuron firing an action potential is explained in terms of how Gibbs free energy is expended to create von Neumann entropy, or a distribution of possible macrostates. This approach therefore models how probabilistic coding in cortical neurons creates a thermodynamic quantity of information. Uncertainty is then resolved during a unitary change of basis, as each cortical neuron, ‘System A’, thermodynamically favors a compatible state with its surrounding environment, ‘System B’. This model of quantum coding is well-established in the literature [17, 18], and is applied here to cortical neurons. This mathematical model of a time-evolving system state, given in terms of matrix mechanics, yields an iterative cycle of information generation and compression – with free energy being physically expended toward information generation and partially recovered upon information compression, in accordance with the Landauer principle [19–22]. At the completion of the computation, as uncertainty is reduced, information is compressed and thermal free energy is released. Due to the relationship between the Gibbs free energy equation and the Nernst equation, the release of free energy shifts the equilibrium potential, leading to sparse but synchronous firing of neurons across the network [23]. The extraction of predictive value reduces entropy, resulting in the selection of a single optimal system state which encodes the state of the surrounding environment.

This report proposes a mathematical model of cortical neuron computation, in which thermal fluctuations materially contribute to signaling outcomes. Additional calculations demonstrate that empirically-grounded estimates for coulomb scattering profiles and decoherence timescales in cortical neurons are consistent with a quantum system, with random electrical noise driving signaling outcomes during a non-deterministic computation.

## II. METHODS

### A. Modeling the cortical neuron as a two-state quantum system

During up-state, cortical neurons linger at their action potential threshold, allowing random electrical noise to prompt a signaling outcome. So, while a neuron is classically interpreted as a binary logic gate in an ‘on’ or ‘off’ state, coded as 1 or 0, it could also be described as having some *probability* of converting to an ‘on’ state or remaining in an ‘off’ state. In this approach, a cortical neuron integrates upstream signals with random electrical noise, defining its voltage state as a function of time, as the system is perturbed. The neuron starts in off-state *ϕ*, not firing an action potential, and over time *t*, it reaches another state *χ*. And so, over some period of time, from *t*_0_ to *t*, the state of the neuron evolves from *ϕ* to *χ*. The timepath taken from one state to another is given by the bra-ket notation:

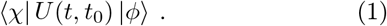

This linear mapping of vectors onto a complex plane provides the probability of a state change, which can be represented in some basis:

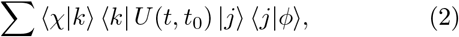

Such that *U* is completely described by base states *k* and *j*:

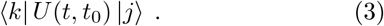

The time interval can be understood as being *t* = *t*_0_ + Δ*t*, so identifying the state of the neuron *χ* at time *t* can be understood as taking a path from one state to another:

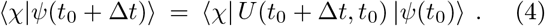

If Δ*t* = 0, there can be no state change. In this case:

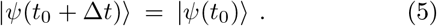

In any other case, the state of the neuron at time *t* is given by the orthonormal base states *k* and *j*, with probability amplitudes:

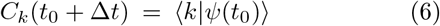

And:

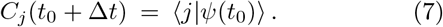

The state vector *ψ* at time *t* is a superposition of the two orthonormal base states *k* and *j*, with the sum of the squared moduli of all probability amplitudes being equal to 1:

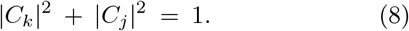

The neuronal state |*ψ*〉 at time *t* can therefore be described as a normed state vector *ψ*, in a superposition of two orthonormal base states *k* and *j*, with probability amplitudes *C_k_* and *C_j_*. With a neuron starting the time evolution in the state |*ψ*(*t*_0_)〉 = *j*, Eq. 4 can be given as:

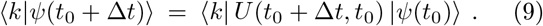

This equation can also be written as the sum of all base states:

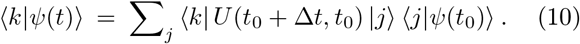

For the state vector *ψ*(*t*), the probability of a state change at time *t* is described by the U-matrix, *U_kj_*(*t*):

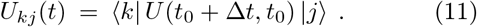

And so, all probability amplitudes are dependent on the amount of time that has passed, Δ*t*:

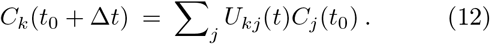

If Δ*t* = 0, there can be no state change and *k* = *j*. If Δ*t* > 0, there is some probability of a state change, where *k* ≠ *j*. As such, the two-state quantum system is described by the Kronecker delta *δ_kj_*:

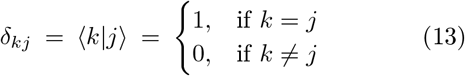

Here, the neuronal state |*ψ_n_*〉 evolves over time, with a binary signaling outcome at time *t* being a function of all local ion states. The state of each ion in the system |*ψ_i_*〉 also evolves with time, with its location at time *t* defined in relation to each local neuron, either inside that neuron or outside it. Any change in the location of a particular sodium ion is a function of the electrochemical potentials of all nearby neurons, which affect the activation state of all local ion channels, and the position, momentum, and energy state of every component electron. If the fundamentally uncertain state of an electron affects the state of the entire ion, in the presence of a dynamic electrical field, the present state of that ion is probabilistic, and the resultant voltage state of each neuron is also probabilistic. This uncertainty is expected to affect the membrane potential of cortical neurons in up-state, with quantum events actually contributing to the probability of a signaling outcome.

### B. Retaining sensitivity to quantum states at the macroscale

Every electron in the system has some possible energy orbital *∊*, and some position in the *x,y,z* plane, both of which are fundamentally uncertain at time *t*. Any energy above ground state may be dissipated toward the production of *information*: a distribution of complex-valued probability amplitudes describing the possible states of each electron. In modeling the electron, the state vector *ψ* can refer to its energy orbital, which can be any one of several orthonormal pure states |*ψ_e_*〉.

Because the state of each electron is fundamentally uncertain, the state of each ion is uncertain. The probabilistic state of each electron affects the most likely state of each ion – and this uncertain state may be sustained in the presence of a dynamic electrical field exerted by each neuron in the vicinity, with the electrical resistance of the neural membrane changing as membrane potentials fluctuate. Because of this sustained uncertainty, each ion can be considered a two-state quantum system, in relation to each neuron. A sodium ion, for example, starts the time evolution outside a given neuron, in state *ϕ_i_*, and has some probability of entering that neuron over time *t*, as it evolves into state *χ_i_*. The location of the sodium ion at time *t*, relative to a given neuron, is represented by the state vector *ψ_i_*, and exists in a superposition of two orthonormal base states with probability amplitudes *C_k_* and *C_j_*.

Because the state of each ion is uncertain, the state of each neuron is uncertain. Since the neuron’s membrane potential is dependent on the location of each ion in the system, and the location of each ion is uncertain, the neuronal membrane potential is also uncertain. A cortical neuron can therefore be considered a two-state quantum system, with some probability of undergoing a state change over time *t*. The cortical neuron starts the time evolution below the threshold for firing an action potential, in state *ϕ_n_*, but has some probability of reaching that threshold over time t, as it evolves into state *χ_n_*. The state of the neuron at time *t*, either having reached the threshold for firing an action potential or not, is represented by the state vector ψn, and exists in a superposition of two orthonormal base states with probability amplitudes *C_k_* and *C_j_*.

In this new approach, either the state of each ion or the state of each neuron can be modeled with the state vector *ψ*. The position of a sodium ion at time *t* has either changed or it has not; the firing state of any neuron at time *t* has either changed or it has not. This model abandons the classical tradition of considering a cortical neuron as a binary computing unit, always in an on-state or an off-state, firing an action potential or not, at any given moment. Instead, it considers the cortical neuron as a two-state quantum system, described by the Kronecker delta. Here, the computational unit calculates the *probability* of firing an action potential, as a function of all probabilistic component states. Since a cortical neuron allows random electrical noise to gate a signaling outcome, and each electron may contribute to the voltage state of multiple neurons, the state of each neuron must be computed simultaneously, as the state of each component pure state is computed. This inherently non-deterministic process can be modeled in mechanistic terms. In this mathematically-grounded approach, the information that is physically generated by a cortical neuron is described as a probability distribution or a mixed sum of all component pure states.

### C. A process of thermodynamic computation in cortical neurons

A cortical neural network is comprised of *N* neurons or computational units, each described by the state vector *ψ_n_*. The state of each neuron |*ψ_x_*〉 at time *t* is uncertain, described in terms of two orthonormal base states. *ρ_x_* is defined as the outer product of this finite dimensional vector space:

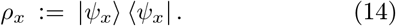

*ρ* is the sum of all mutually orthogonal pure states *ρ_x_*,each occurring with some probability *p_x_*:

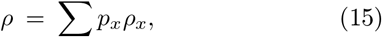

The statistical ensemble of these component pure states is the physical quantity of information held by the system, given by the von Neumann entropy formula [24]. This thermodynamical and computational quantity is a high-dimensional volume of probability, represented by the trace across the density matrix *ρ*:

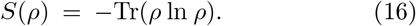

The effect of integrating uncertainty across mixed component states is to increase dimensionality and information simultaneously. Here it is useful to consider an ensemble of neurons, each with a state vector *ψ_x_*. The mixed sum of all component states is:

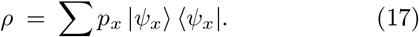

The expectation value 〈A〉 of the observable *A* is given by:

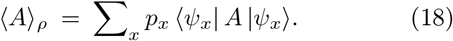

An observation *A* can then be made by measuring the system state *ρ*:

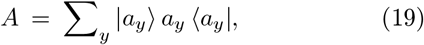

In a mixed state, any one outcome *a_y_* occurs with probability *p*(*a_y_*):

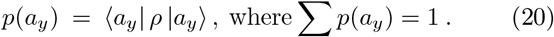

The quantity of Shannon entropy *H*(*y*) for the ensemble of measurement outcomes {*a_y_*, *p*(*a_y_*)} is always greater than the quantity of von Neumann entropy *S*(*ρ*), reaching equality only if *A* and *ρ* perfectly commute:

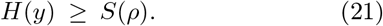

This is because the number of possible states for the system (an algebraic quantity) may be greater than the dimensionality needed to represent any linearly correlated states (a geometric quantity). Only the noncorrelated states, which do not commute, will yield an uncompressible density matrix. And so, the randomness of information is *minimized* upon measurement if the measured observable commutes with the density matrix, leading to a more predictable value being measured.

By contrast, the quantity of entropy is *maximized* when the system state is completely random and all non-zero eigenvalues have equal probability *p_x_*. In general, the von Neumann entropy of the system is less than maximal when some system states are more likely than others [25, 26]. This will always be the case with a biological system, which does not have a completely random system state. Given that *ρ* has *D* non-vanishing eigenvalues, the value of *S*(*ρ*) is always less than or equal to the logarithm of that number:

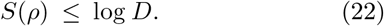

The entropy is also maximized when less is known about the previous system state. For all non-vanishing eigenvalues *p*_1_, *p*_2_,…*p_n_* ≥ 0, which together equal a normalized probability distributionp *p*_1_ + *p*_2_ + … +*p_n_* = 1, the entropy for the system is greater if the previous system state is not known:

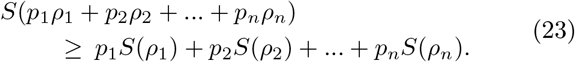

As the system is *perturbed*, by interacting with its environment, the distinguishability between states may be lost with the addition of this new information. This applies to the state of each ion, or the state of each neuron, as it interacts with its surrounding environment. If non-orthogonal pure states are mixed, then *S*(*ρ*) < *H*(*y*). If all component pure states are completely orthogonal, then *S*(*ρ*) = *H*(*y*). Therefore, entropy is *additive* in uncorrelated systems, but it is *subtractive* in correlated systems, as redundancies or correlations between system states are reduced. This leads to the subadditivity rule:

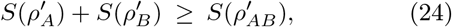

Which applies to any ion, any neuron, or the entire cortical neural network: an open non-equilibrium thermodynamic system (e.g. System A), comprised of *N* units, each described by a state vector *ψ_Ax_*, operating within an environment (e.g. System B), comprised of *M* units, each with a state vector *ψ_Bx_*. Each system is described by a density matrix, or a mixed sum of component pure states. The system and its surrounding environment are initially uncorrelated with each other, and the combined system is created by the tensor product of the two density matrices:

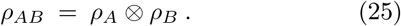

The combined system evolves over time, as System A is perturbed by System B:

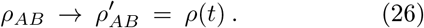

If the systems are uncorrelated, the states are additive and the entropy remains unchanged:

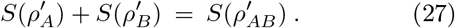

But if any states are correlated, or non-distinguishable, entropy will be compressed:

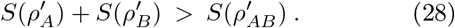

As System A is perturbed by System B, the combined system evolves over time, with the combined density matrix undergoing a unitary change of basis:

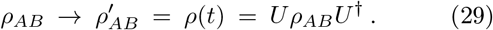

The unitary change in basis is provided by the time shift operator *U*:

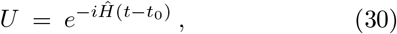

Where the natural log of *U* is equal to the imaginary unit *i* (which generates a complex axis), multiplied by the Hamiltonian operator 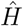 (the sum of all potential and kinetic energies in the system, regardless of whether that system is an ion, a neuron, or the entire neural network), and the amount of time that has passed (*t* − *t*_0_), over the reduced Planck constant 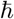. *U* is represented by a square matrix of finite dimensionality:

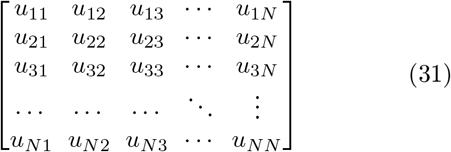

The product of this *U*-matrix and its complex conjugate *U*^†^ is an identity whose determinant is always equal to 1:

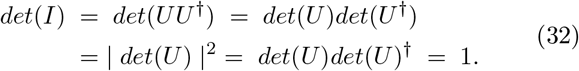

The columns of U form an orthonormal set of column matrices, and the rows form an orthonormal set of row matrices. If System A and System B become correlated during the unitary change in basis, the rows and columns of the matrix become linearly dependent vectors. This internal consistency allows compression of the combined matrix. The reduction in von Neumann entropy yields observables, or eigenvalues, on the boundary region of the probability density, yielding an observable system state. That is, if time and energy commute, energy is redistributed to realize an optimal system state from the distribution of possible system states. But during the computational process, both the amount of time that has passed and the system state remain undefined. If ‘System A’ and its surrounding environment ‘System B’ become correlated over the course of this probabilistic time evolution, then information compression or entropy reduction will occur as two systems interact:

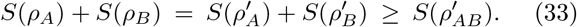

This equation reaches equality if all component pure states remain completely uncorrelated. If the two systems become correlated, then the information (or distinguishable states) that were initially encoded in the separate systems are now encoded in the quantum entanglement of the combined system. The entanglement is given as the tensor product of two Hilbert spaces 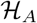 and 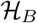:

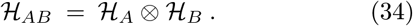

Hilbert spaces are inner product spaces with an orthonormal basis, populated by probability densities, which are generated by linear transformations of vectors performed by linear operations. Each orthonormal pure state generates a Hilbert space, and any pure states that are identical cannot physically co-exist. Identifying non-distinguishable states (or linear correlations between pure states), through a unitary change of basis, will compress the von Neumann entropy of the combined system.

### D. Schmidt decomposition and Schumacher compression

The combined system, an M × N density matrix, undergoes eigendecomposition through a unitary change of basis. The eigenvalues, given by *χ*, are identified during a series of linear operations, given by the M × M matrix *U*, some measurement or observation *d*, and the N × N matrix *V*^†^:

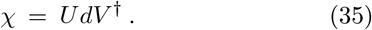

A mixed sum of orthogonal pure states, given by density matrix *ρ*, has multiple possible realizations, for ex-ample *ρ*_1_:

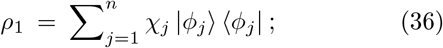

And *ρ*_2_:

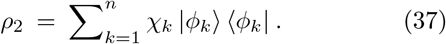

Here, |*ϕ_j_*〉 and |*ϕ_k_*〉 are bases within the Hilbert spaces 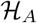 and 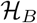, respectively. *χ_j_* and *χ_k_* are eigenvalues within the set of eigenstates *χ_i_*. All values for *χ_i_* are positive and normalized, such that:

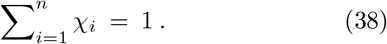

Here, *n* is the minimum number of possible states represented in Hilbert space, and the square root of *χ_i_* is the Schmidt coefficient, given by the partial trace across the two density matrices. Any normalized separable states form a convex set, with every bipartite pure state 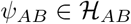 within the combined system A ∪ B [27]. Two different measurements lead to two different state purifications, yielding two different realizations. Two possible purifications |*ψ_AB_*〉 for a bipartite pure state are given by 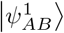 and 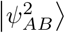. These are possible outcomes of the interaction between System A and System B:

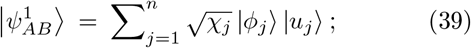

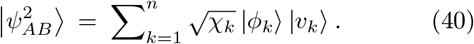

The two purifications only differ by a unitary transformation acting on the combined system, System AB. Therefore, there must be some unitary matrix *d* which converts one state to another:

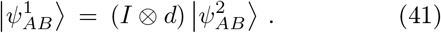

Different ensemble realizations of a density matrix can therefore be identified simply by making different measurements, or observations, given by *d*:

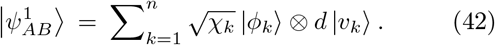

This method, called Schmidt decomposition, permits quantification of the separability between two systems [28, 29]. Each state |*ψ_AB_*〉 is separable if and only if there is one non-zero Schmidt coefficient. If more than one Schmidt coefficient is non-zero, the state is entangled and represented redundantly by both systems. If all Schmidt coefficients are non-zero, the systems are maximally entangled. Schmidt decomposition thereby allows quantification of the entanglement of a bipartite system [30, 31]. In this case, an ion, a neuron, or the entire network is the open non-equilibrium ‘System A’, with *N* units, operating in surrounding ‘System B’, with *M* units. This method usefully allows quantification of the entanglement *∊* for a multipartite system of *R* components, with *n* possible states [32]:

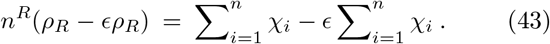

The compression of information entropy that occurs over a time evolution in an open thermodynamic system, as the system state becomes correlated with the state of its environment, is an entirely natural process of information compression. A system of *N* units has some quantity of information entropy, given by the mixed sum of all component pure states, after some time evolution *t*. These component pure states are described by normed state vectors in a Hilbert space, and they yield a two-state quantum system, with each state vector evolving from state *ϕ* to state *χ* over time *t*. As System A interacts with its surrounding System B, any linear correlations reduce the number of distinguishable pure states in the combined system. This reduction of quantum information is proportional to the corresponding reduction in Hilbert space:

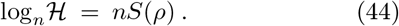

This relation is the Schumacher equation for the compressibility of an ensemble of orthonormal pure states. It is predicted to apply to any non-equilibrium ‘System A’ within an environmental ‘System B’. If System A acts as a net energy sink, while retaining a stable overall temperature, System A may use this free energy to drive computational work. Here, the state of each electron, each ion, or each neuron can be described as the following set of pure states:

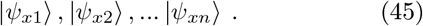

Each pure state occupies a Hilbert space 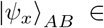, with an ensemble given by { |*ψ_x_*〉_*AB*_, *p_x_*}. The density matrix representing all mutually orthogonal pure states is therefore defined as an outer product:

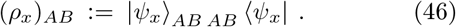

A Schumacher compression allows System B to be discarded as correlated states are identified, by taking a partial trace across the combined density matrix [18, 33]. At the start of the computation, the density matrix for the combined ensemble of mixed states is given by all state vectors across the neural network, provided by the sum of all pure states *ρ_x_*, each occurring with probability *p_x_*:

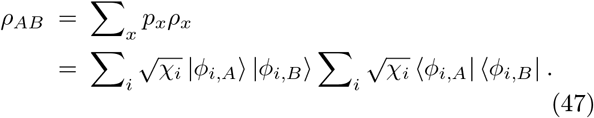

A pure state shared between System A and System B can then be realized through a purification:

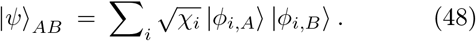

Any pure state that is shared between System A and System B will be realized by taking the partial trace across the combined density matrix. This process demonstrates that a single realization of pure states can be represented *independently* by the reduced density matrix of either system:

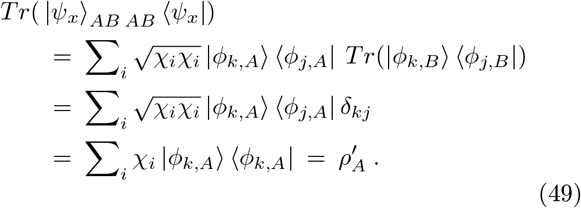

Since 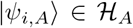 and 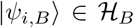 are orthonormal sets, then *χ_i_* with (*i* = 1, 2,…*N_s_*) are the eigenvalues of both *ρ_A_* and *ρ_B_*. If the two density matrices have the same spectrum of eigenvalues, these redundancies can be reduced or compressed. This allows System A to take on an actualized state by realizing correlations with System B. Taking the partial trace of *ρ_AB_* over B yields the mixed state of subsystem A:

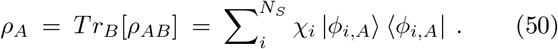

And taking the partial trace of *ρ_AB_* over A yields the mixed state of subsystem B:

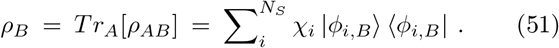

Block diagonalization of the matrix then realizes the eigenvalues from this ensemble of mixed states:

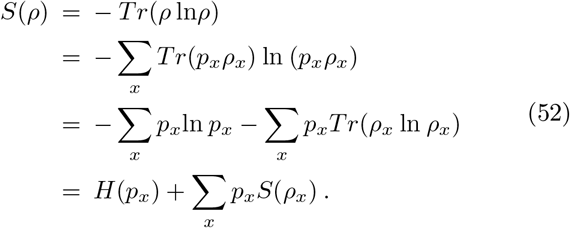

This value, *H*(*p_x_*), is the Holevo bound for the quantity of information in the ensemble {*ρ_x_*, *ρ_x_*} with trace *ρ_x_* = 1 for each *x* [17, 34]. As *ρ* is realized for an ensemble of mutually orthogonal mixed states during a computation, the overall quantity of information held by the system is compressed:

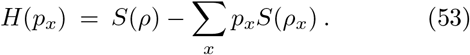

Due to the concavity of the log function, the value of *H*(*p_x_*) is always positive [35, 36]. Therefore, by geometric necessity, the total quantity of von Neumann entropy generated is always greater than the quantity of compression, respecting the second law of thermodynamics.

## III. RESULTS

### A. The extraction of predictive value is equivalent to a process of quantum information compression

The compression of information is a thermodynamic process that generates a discrete quantity of free energy [19–22]. The conversion of these quantities is defined by the Landauer limit, which calculates the amount of free energy released by the erasure of a single digital bit:

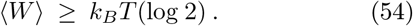

This concept can be extrapolated to non-digital computational operations with any number of possible system states. The recently-derived Still dissipation theorem demonstrates the quantity of non-predictive information in a system is proportional to the energy lost when an external driving signal changes by an incremental amount within a non-equilibrium system [37]. Here, the amount of information without predictive value is equivalent to the amount of work dissipated to entropy (*W*_diss_) as the signal changes over a time course from *x*_*t*=0_ (the immediately previous system state) to *x_t_* (the present state):

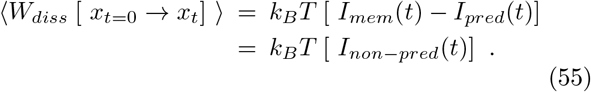

This equation stipulates that any information stored in memory, subtracted by the total amount of information with predictive value, is the amount of non-predictive information remaining after predictive value has been extracted. This quantity is dissipated as ‘work’ upon the completion of a computation, and therefore is unavailable as free energy to accomplish work within the system [38]. The memory held by the system *I*_mem_(*t*) is the amount of entropy gained as System A interacts with System B, given by *S*(*ρ*). The predictive value *I*_pred_(*t*) is the amount of compression achieved by reducing correlated states during the interaction 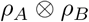, and the net quantity of entropy remaining *I*_non-pred_(*t*) is the amount of work dissipated by the system after completing one full thermodynamic computation:

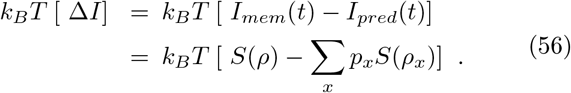

*I*_mem_(*t*) − *I*_pred_(*t*) ≥ 0, otherwise the system would create energy through information processing alone, an event forbidden by the first law of thermodynamics. Likewise, the quantity of Holevo information *H*(*p_x_*) was shown in Eq. 53 to be a positive quantity by geometric necessity; this value must be positive to prevent a similar violation of the first law of thermodynamics. Yet the smaller the value of Δ*I* = *I*_mem_(*t*) − *I*_pred_(*t*) = *H*(*p*_x_), the more compression has occurred during a computational cycle. This event minimizes net entropy production and maximizes free energy availability. During this thermodynamic computation, predictive value is extracted, the entropy or number of possible system states is reduced, and the system becomes more ordered. As correlations are identified between the two systems, the most consistent system state is selected from a distribution of possible system macrostates, and information is compressed:

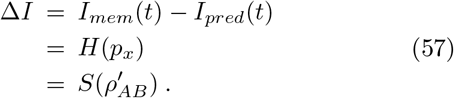

Δ*I* is always greater than or equal to zero, even though disorder is reduced during the computation, as the optimal system state is identified. To avoid creating energy during the computation – a prohibited outcome – the quantity of Δ*I* must greater than or equal to zero, with any non-predictive information, *I*_non-pred_(*t*), being held in memory until predictive value can be assigned:

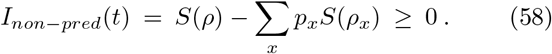

The amount of predictive value extracted, *I*_pred_(*t*), is positive if linear correlations can be identified in the combined density matrix during the computation:

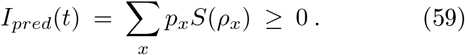

But this quantity of predictive value can be no more than the starting quantity of entropy, *I*_mem_(*t*), from which predictive value is extracted:

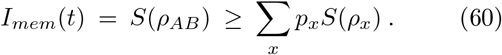

A thermodynamic computation involves an extraction of predictive value, a compression of information entropy, and a reduction in possible system states. These thermodynamic and computational processes are equivalent:

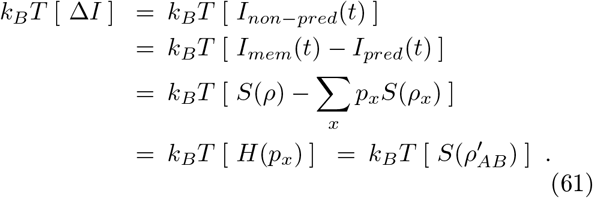

That is: the quantity of information with predictive value is equivalent to the amount of information compression that occurs during a computational cycle, and that quantity places a thermodynamic limit on the amount of increased order achieved by the system. Although entropy is generated by the system, predictive value can be extracted, leading to a reduction in the distribution of possible system states. The compression of entropy is paired with a release of free energy, in accordance with the Landauer principle [19–22]. And so, during the computational process, random electrical noise is converted to *work*, with any reduction of uncertainty materially contributing to the probability of a signaling outcome.

### B. A cortical neural network demonstrably meets the criteria for a quantum system

If biological neural networks perform quantum computations, then entire atoms within these thermodynamic systems must be shown to sustain coherent states, and these coherent states must contribute to the selection of an optimal system state from a probability distribution. Yet doubt has been cast on this hypothesis. Twenty years ago, Max Tegmark published a notable paper explaining why “the brain is probably *not* a quantum computer” [41]. The report convincingly argued the brain must be a classical system, because rates of quantum decoherence do not occur on the timescales relevant to neural activity, namely the time between action potentials (0.05 - 200 spikes per second) or the time taken to spike (approximately 1 ms). These timescales rely on estimates of coulomb scattering of sodium ions at the neural membrane. The original calculations are worth revisiting, particularly since recent studies have updated the estimates for coulomb scattering rates at the neuronal membrane.

It is provided in the Tegmark paper (Section 3.2.1) that Λ, the scattering rate, should be defined as Λ ≡ *n σ υ*, “where *n* is the density of scatterers, *σ* is the scattering cross-section, and *υ* is the velocity”.

This definition of the scattering rate Λ is not debated. It is also provided in the paper: “The spatial superposition of an ion decays exponentially on a time scale Λ^-1^ of order its mean free time between collisions. Since the superposition of the neuron states ‘resting’ and ‘firing’ involves *N* such superposed ions, it thus gets destroyed on a timescale *τ* ≡ (*N*Λ)^-1^.”

This definition given for the decoherence timescale *τ*_dec_ is also not debated. However it must be pointed out that a neuronal state change is commonly achieved with a change in position of far fewer ions than was estimated in the original paper - particularly in cortical up-states, where neurons remain near the threshold for firing an action potential [42]. Given this agreement that:

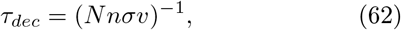

the values of *N*, *n*, *σ* and *υ*, which together delimit the timescales of quantum decoherence, can now be formally reassessed.

The density of Na+ ions is given by 120 mM or *n* = 7.23 × 10^16^ m^-3^. Meanwhile, the velocity of each Na+ ion at 37°C (310K) can be estimated as *υ* ≈ sqrt (*k*_B_*T*/*m*) = 10.6 m/s. Both of these estimates are identical to those given in the original paper.

Now, it is worth more closely considering the crosssectional area σ. The scattering profile of Na+ ions at the lipid membrane has been studied in greater detail since these initial estimates of ion decoherence in biological systems were published [43, 44]. The new scattering estimates demonstrate ion-lipid membrane interactions to reach equilibrium over 100 - 120 ns in a system with 200 mM NaCl extracellular concentration and 20 mM KCl intracellular concentration. While these studies simulate ion kinetics at the lipid membrane of a generic cell, the ion density and lipid membrane properties render this estimate highly relevant to the neural membrane. This model gives a scattering cross-sectional area of *σ*=32.8 nm^2^ or 3.28 × 10^-17^ m^2^ across the lipid membrane, with a mean residence time of 465 ps for a single Na+ complexing with the lipid membrane [43].

It should be noted that any interactions between Na+ ions in the extracellular milieu do not alter the state of the neuron. The decoherence, or resolution of the neuronal state, only occurs when these ions interact with the neuronal membrane. We can estimate that approximately N = 100 Na+ ions must cross a neuronal membrane to induce a state change, a reasonable assumption in the case of a cortical up-state, when neurons are already approaching action potential threshold.

Together, these estimated values provide a decoherence timescale *τ*_dec_of 0.4 ms:

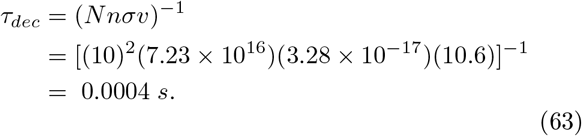

This value provides the estimated rate at which neurons resolve their voltage state, thereby defining the temporal parameters for the summation of coincident charge flux across the neuronal membrane. This decoherence rate of 2500 Hz is similar to the exchange rate of Na+K+ pumps embedded in the neuronal membrane, which pump sodium ions out of the cell at a variable frequency of 1000 Hz to 100,000 Hz [45, 46]. The rate of stochastic charge flux is essentially balanced by this rate of sodium-potassium exchange. And so, any burst in temporally-coincident events within this timeframe will affect the probability of a neuron firing. As a result, the compression of information (paired with free energy release) is expected to cause fluctuations in the neuronal membrane potential, shifting the likelihood of the neuron reaching action potential threshold and opening voltagegated ion channels.

The timescales of decoherence are key, particularly in relation to the estimated timescales of ion dynamics and ion dissipation. From the Tegmark paper: “If *τ*_dyn_ ≪ *τ*_dec_, we are dealing with a true quantum system, since its superpositions can persist long enough to be dynamically important. If *τ*_dyn_ ≫ *τ*_diss_, it is hardly meaningful to view it as an independent system at all, since its internal forces are so weak that they are dwarfed by the effects of the surroundings. In the intermediate case where *τ*_dec_ ≪ *τ*_dyn_ < *τ*_diss_, we have a familiar classical system.”

A comparison of Na+ ion dynamics, dissociation, and decoherence at the neuronal membrane shows that *τ*_dyn_ ≪ *τ*_dec_ and *τ*_dyn_ < *τ*_diss_. The timescales of Na+ ionization dynamics τ_dyn_are on the order of 0.5 to 5 ps [47], and the timescales of ion dissociation from the lipid membrane *τ*_diss_ are on the order of 400 to 600 ps [43]. Na+ ions readily displace each other at the membrane, and so the collision timescale between Na+ ions and lipid macromolecules *τ*_coll_ is equivalent to the cited mean residence time of 465 ps. In the Tegmark paper, the dissipation rate is given by the time between collisions, multiplied by the sodium ion mass and divided by the mass of the water molecule:

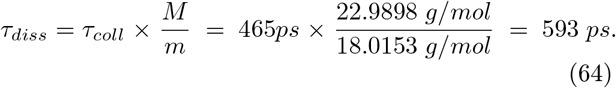

These timescales of sodium ion dissipation *τ*_diss_ and sodium ionization dynamics *τ*_dyn_, respectively, are approximately one million and one hundred million times shorter than the newly estimated timescales of decoherence between sodium ions and the neural membrane, calculated here to be on the order of 0.4 ms. Together, these timescales fit the Tegmark criteria for a quantum system. Yet interestingly, it should be noted that only neurons maintaining a cortical up-state will be able to participate in these quantum computations, as the quantitative requirements for ion flux are too high at normal resting potential to sustain a coherent state (N > 10^6^ Na+ ions). As such, peripheral neurons and subcortical neurons are expected to behave deterministically, rather than probabilistically. Only neural systems which hover near the threshold for a state change should sustainably generate quantum information. Similarly, the calculation of eigenvectors, by a unitary change of basis, should contribute to signaling outcomes only in neural systems which reside near the threshold for a state change.

In summary, cortical neural network computation is expected to be vindicated as room-temperature quantum computing, achieved by calculating eigenvalues for the position, momentum, and energy state of each electron in the system, after some perturbation has occurred. The above calculations demonstrate that interactions between Na+ ions and the lipid membrane, based on mass, velocity, ion density, collision rate, and reasonable estimates of coulomb scattering yield a decoherence rate of 0.4ms, which is much longer than originally estimated. This timeframe is approximately ten times shorter than the minimum time between spikes, which is 5 ms in fastspiking cortical neurons. That difference is expected to correlate with the minimum time needed to overcome hyperpolarization and re-establish the electrochemical resting potential to a cortical up-state. Given these newly estimated timescales of sodium ion decoherence, dissipation rates and ionization dynamics, cortical neural networks can reasonably be theorized to have the properties of a quantum system. It is therefore expected that quantum information processing contributes to thermal fluctuation-dissipation dynamics, thereby affecting the electrical resistance of each computational unit.

### C. Specific predictions of this model

If cortical neural networks are indeed quantum computing systems rather than classical computing systems, then evidence of quantum information generation and compression should be observed in the neocortex. As such, this theory makes specific predictions, with regard to the wavelength of thermal free energy release upon information compression [48], as well as the expected effects of electromagnetic stimulation and pharmacological interventions in cortical neural networks [49]. Some additional predictions of the theoretical framework, prompted by the present model, include:

#### 1. Predicted coulomb scattering rates of sodium ions at the neuronal membrane

Coulomb scattering of sodium ions at the cellular membrane should be *σ* ≈ 32.8nm^2^ in cortical neurons. Given the known density of sodium ions in the extracellular milieu and effects of the dynamical electrochemical potential of the cortical neuron membrane in vivo, the scattering rate of sodium ions should be experimentally observed to be in the range of tens of square nanometers.

#### 2. Predicted decoherence timescales of sodium ions at the neuronal membrane

Timescales of sodium ion decoherence at the cellular membrane should be *τ*_dec_ ≈ 0.0004s in cortical neurons. Given the known density of sodium ions in the extracellular milieu, the high sensitivity of cortical neurons to stochastic charge flux, the velocity of sodium ions at normal body temperature, and the dynamical electrochemical potential of the cortical neuron membrane, timescales of sodium ion decoherence should be longer than timescales of ionization dynamics and timescales of thermal dissipation (*τ_dyn_* < *τ_diSS_* ≪ *τ_dec_*).

#### 3. Predicted ionization and dissociation dynamics of sodium ions at the neuronal membrane

Timescales of Na+ ionization dynamics should be empirically shown to be on the order of 0.5 to 5 ps [47], and the timescales of ion dissociation from the lipid membrane should be empirically shown to be on the order of 400 to 600 ps [43]. If these timescales of sodium ion dissipation *τ*_diss_ and sodium ionization dynamics *τ*_dyn_, respectively, are much shorter than the timescales of decoherence between sodium ions and the neural membrane, then the system meets the criteria of a quantum system.

#### 4. Van der Waals forces preceding the action potential

Van der Waals forces should occur between extracellular sodium ions and lipid molecules within the neural membrane. This prediction may be tested with an atomic force microscope, as cortical neurons reach action potential threshold (or don’t) on the basis of apparently stochastic membrane potential fluctuations and random electrical noise. In cortical neurons, spontaneous changes in atomic proximity should cause measurable van der Waals forces and increased sodium ion leak across the cellular membrane ahead of signaling events. In peripheral neurons, by contrast, the movement of sodium ions should be consistently random.

#### 5. Random electrical noise affects signaling outcomes in individual cortical neurons

Mathematical models of probabilistic neural coding that incorporate random noise have proven useful in predicting cortical activity at both the cellular level and the systems level [51, 52]. And yet, the mechanisms by which cortical neurons regularly select a statistically unlikely but advantageous system state, in the context of a noisy dataset, are not well-understood [5, 6, 52]. The most optimal system state must somehow be selected from a probability distribution, in accordance with thermodynamical laws. Here, it is predicted that the non-deterministic signaling outcomes of cortical neurons result from the compression of entropy, as predictive value is extracted. If quantum information processing does contribute to signaling outcomes in cortical neural networks, then taking a Fourier transform of electrical noise should yield an improved prediction of the inter-spike interval, compared with models that only sum upstream inputs and work effectively for modeling spinal reflex circuits. If signaling outcomes in cortical neurons are truly deterministic, then random electrical noise should have no effect on the timing of action potentials, and classical mechanisms should fully explain the highly variable inter-spike intervals that are observed *in vivo*.

#### 6. Random electrical noise affects network-level dynamics

Information compression events should cause synchronized neural activity in sparsely-distributed populations across neocortex. The state of the neuron at the moment the Hamiltonian operator is resolved, the density matrices commute, or the wavefunction collapses, will determine its response. Neurons in a cortical up-state, allowing inherently stochastic events to gate a signaling outcome, may be nudged toward (or away from) action potential threshold by the local redistribution of energy, as the Hamiltonian is resolved. Therefore, a systemwide computational event, resolving the uncertainty in all component pure states, should resolve all membrane potentials and lead to synchronous neural activity across the network. This synchronous activity should be evident only in neural circuits that allow random electrical noise to gate signaling outcomes, and this synchronous activity should occur periodically, as information is physically generated and compressed.

## IV. DISCUSSION

Modeling a unitary change of basis by an ion, a neuron, or the entire brain, ‘System A’ – in the context of its surrounding environment, ‘System B’ – results in the selection of an optimal system state from a large probability distribution, through an inherently probabilistic computational process that is described in terms of matrix mechanics. Once information compression occurs, and the optimal system state for that ion, that neuron, or that brain is selected within its immediate surrounding environment, a new time evolution begins, and a new distribution of probabilistic macrostates is generated, in accordance with the von Neumann projection postulate. This process of information generation and compression then repeats, yielding iterative computational cycles that allow a thermodynamic computing system to gradually improve in predictive capacity.

If the system successfully extracts predictive value, the quantity of von Neumann entropy decreases, and the system can achieve a more ordered state during a computational cycle. If the system becomes more disordered, with a net increase in the distribution of possible system states, there is scope for extracting predictive value and undergoing a useful compression event. Interestingly, both things can happen within a single computational cycle. But the net value of *W_diss_* = *k_B_T*(Δ*I*) = *T*Δ*S* must always be greater than or equal to zero. If this quantity were negative, more predictive value would be extracted from the information than the total amount of information available. The first law of thermodynamics holds, and energy simply cannot be created during the computation. A strict accounting also requires that free energy must be physically released back into the system during information compression, in accordance with the Landauer principle. This event is expected to be paired with a resolution of the system state, with non-deterministic outcomes being actualized across the neural network - an event characterized by sparse yet synchronous activity across the neural network.

It is worth noting the deep connection between information processing and thermodynamic efficiency. The amount of Shannon entropy generated, as an internal process learns about an external process, is bounded by the total quantity of thermodynamic entropy [39]. In other words, the rate at which non-compressible information is generated provides a lower bound on temperature-normalized energy consumption [40]. There is a direct link here between energy efficiency and compressibility, which is ultimately dependent on the consistency or predictive value that can be extracted from a thermodynamic quantity of information.

It is therefore also worth noting the deep connection between energy-efficient computation and predictive processing. Predictive power is maximized in non-Markovian multi-layer networks through oscillatory kernels – and in particularly noisy systems, the system is not memoryless, but rather capitalizes on signal values from the past that are relevant in assessing the present and predicting the future [53]. The system under consideration – a cortical neural network engaging in probabilistic coding – is not Markovian, with a stationary transition matrix. It is a continuous stochastic process, with any state change prompted by an exponential random variable: the U-matrix. This system has a finite or countable state space, with dimensions equal to the transition matrix; it has some initial state *j*; and it has some non-negative number of computational units *N*, each having some probability of transitioning from *j* to *k*. The posterior probability, at the completion of a computation, is dependent on the state of the environment, the amount of time that has passed, and the history of system states. It is not independent of its environment, it is not independent of the amount of time that has passed; and it is not memoryless, as in a Markovian process.

To quantify the amount of time that has passed, Δ*t*, as work is dissipated toward non-predictive information over the course of a single computational cycle, a semi-Markovian model of the Still dissipation theorem is specifically employed [40]. Here, Δ*t* is the timescale over which “an external driving signal changes by an incremental amount, thereby doing work on the system” [37]. This semi-Markovian process transitions the system state from a prior probability (the ‘past’) to a posterior probability (the ‘present’) through a time-dependent unitary change of basis, thereby achieving Bayesian inference. The total amount of work dissipated to non-predictive information over the course of a computation is equivalent to *I*_mem_ (the total quantity of information gained during the computation) minus *I*_pred_ (the predictive value extracted from that quantity). This entire process of information generation and compression occurs over Δ*t*, and therefore this net quantity is summed over the entire protocol, in a non-reversible computational process.

During this non-reversible computational process, information is compressed and free energy is released back into the system. This thermal free energy release, at synapses whose uncertainty is reduced, is paired with the realization of a single system macrostate from some distribution of possible system macrostates. Usefully, this model of thermodynamic computation directly relates the energy-efficiency of the encoding system to the extraction of predictive value.

In cortical neural networks, an optimal system state in the present context must be selected from a large probability distribution. Neuroscientists have previously modeled this inherently probabilistic computation with Bayesian statistics [50], random-connection models [51], or fanofactor analysis of spike variance over time [54]. This report shows that non-deterministic signaling outcomes can be achieved through a mechanistic (not a statistically random) process. Here, cortical neurons physically generate information through noisy coding, then compress that information to achieve non-deterministic signaling outcomes.

In this model, a system-wide computation is achieved by a cortical neural network through the cyclical generation and compression of von Neumann entropy. The macrostate of the system is resolved as every component pure state is resolved. This process periodically culminates in a defined system state, with a non-deterministic outcome for every computational unit, as predictive value is extracted. Due to the utility of this approach, it may be reasonable to consider cortical neurons as qubits, which calculate the probability of reaching a state change, rather than classical computing units, which exist in a simple on-or-off state.

## ACKNOWLEDGMENTS

The author received support for this work from the Western Institute for Advanced Study, with generous donations from Jason Palmer, Bala Parthasarathy, and Vanguard Charitable.

